# The *Candida* Genome Database: Annotation and Visualization Updates

**DOI:** 10.1101/2024.10.01.616131

**Authors:** Jodi Lew-Smith, Jonathan Binkley, Gavin Sherlock

## Abstract

The *Candida* Genome Database (CGD; www.candidagenome.org) is unique in being both a model organism database and a fungal pathogen database. As a fungal pathogen database, CGD hosts locus pages for five species of the best-studied pathogenic fungi in the *Candida* group. As a model organism database, the species *Candida albicans* serves as a model both for other *Candida* spp. and for non-*Candida* fungi that form biofilms and undergo routine morphogenic switching from the planktonic form to the filamentous form, which is not done by other model yeasts. As pathogenic *Candida* species have become increasingly drug resistant, the high lethality of invasive candidiasis in immunocompromised people is increasingly alarming. There is a pressing need for additional research into basic *Candida* biology, epidemiology and phylogeny, and potential new antifungals. CGD serves the needs of this diverse research community by curating the entire gene-based *Candida* experimental literature as it is published, extracting, organizing and standardizing gene annotations. Most recently, we have begun linking clinical data on disease to relevant Literature Topics to improve searchability for clinical researchers. Because CGD curates for multiple species and most research focuses on aspects related to pathogenicity, we focus our curation efforts on assigning Literature Topic tags, collecting detailed mutant phenotype data, and assigning controlled Gene Ontology terms with accompanying evidence codes. Our Summary pages for each feature include the primary name and all aliases for that locus, a description of the gene and/or gene product, detailed ortholog information with links, a JBrowse window with a visual view of the gene on its chromosome, summarized phenotype, Gene Ontology, and sequence information, references cited on the summary page itself, and any locus notes. The database serves as a community hub, where we link to various types of reference material of relevance to *Candida* researchers, including colleague information, news, and notice of upcoming meetings. We routinely survey the community to learn how the field is evolving and how needs may have changed. A key future challenge is management of the flood of high-throughput expression data to make it as useful as possible to as many researchers as possible. The central challenge for any community database is to turn data into knowledge, which the community can access, use, and build upon.

## Introduction

The *Candida* Genome Database annotates research data for five species within the *Candida* genus, and will likely add others in the future. These species are *C. albicans, C. glabrata, C. dubliniensis, C. parapsilosis*, and *C. auris*. Among these, *Candida albicans* is by far the best studied of these fungal pathogens and serves as a model organism for the study of less experimentally tractable members of the group (Kabir *et al*. 2012a).

*Candida albicans* causes two types of disease in humans. The less lethal candida diseases are common among both immunocompetent and immunocompromised patients and include dental caries and mucosal candidiasis of the oral cavity and the genitourinary tract (Pfaller and Diekema 2007; Schelenz 2008; Giri and Kindo 2012; Eidt *et al*. 2020; Vila *et al*. 2020; Lass-Florl *et al*. 2024). However, in immunocompromised patients, invasive infection of the bloodstream, abdominal organs, and central nervous system can lead to mortality rates exceeding 35% (Fisher-Hoch and Hutwagner 1995; Groll *et al*. 1998; D’ENFERT 2009; Pappas *et al*. 2018; Lass-Florl *et al*. 2024). Indeed, *Candida albicans* is the third or fourth most common nosocomial bloodstream isolate and treatment is costly and significantly extends the length of hospitalization (Calderone 2002; Ostrosky-Zeichner and PAPPAS 2006; Pfaller and Diekema 2007; D’enfert 2009; Andes *et al*. 2012; Dodds ASHLEY *et al*. 2012).

While at least 15 distinct *Candida* spp. have been shown to cause human disease, the majority of invasive infections are caused by *Candida albicans, C. glabrata, C. parapsilosis, C. tropicalis, C. krusei*, and, more recently, *C. auris* (Kullberg and Arendrup 2015; Antinori *et al*. 2016; Mccarty and Pappas 2016; Lamoth *et al*. 2018; Mccarty *et al*. 2021). Within the past ten years, *C. auris* has become a significant global concern on almost every continent (Rhodes 2019), causing large outbreaks in both pediatric and adult populations within healthcare settings (Rhodes 2019; Hui *et al*. 2024; Lass-Florl *et al*. 2024; Sokou *et al*. 2024). Drug resistance is especially problematic for *C. auris*, resulting in the U.S. Centers for Disease Control recently classifying this fungus as an urgent antimicrobial-resistance threat due to its common resistance to multiple antifungal drugs, ability to spread easily via skin infection in nosocomial settings, and the severity of infections (Cdc 2023).

More basic research into non-albicans species has become necessary, as these species have distinct pathogenic profiles. For example, fluconazole resistance is a concern for *C. glabrata, C. parapsilosis*, and *C. auris*, but less so in *C. tropicalis* infection (Lass-Florl *et al*. 2024). Where *C. parapsilosis* most commonly infects neonates and infants (Chow *et al*. 2012; Toth *et al*. 2019), *C. glabrata* is more frequently isolated in older patients and thus has a higher geographical distribution in high-income countries and regions with aging populations (Pfaller *et al*. 2019; Lass-Florl *et al*. 2024). An intriguing aspect of *Candida* biology is that some species more commonly undergo filamentation and form biofilms, while others more often remain in the planktonic yeast forms and yet cause severe disease in mammals.

*C. albicans* was formerly classified in the same order as *Saccharomyces cerevisiae*, but has recently been reclassified to the order *Serinales* (Chen *et al*. 1984; Hendriks *et al*. 1991; Groenewald *et al*. 2023) to reflect higher genome-informed diversity within the subphylum *Saccharomycotina. S. cerevisiae* and *C. albicans* are separated by ∼300 million years of evolution (Hedges *et al*. 2015; Vlastaridis *et al*. 2017). Despite sharing many similarities, including 3,453 pairs of orthologous genes, the two fungi inhabit very different environmental niches, with *S. cerevisiae* existing as a saprophyte and *C. albicans* living in close association with its mammalian hosts. Since their last common ancestor, extensive “transcriptional rewiring” has occurred in their regulatory networks. Further, *C. albicans* exists almost exclusively as a diploid organism, while *S. cerevisiae* lab strains are typically haploid, though diploidy occurs commonly in the wild and is required during the sexual cycle.

Because *C. albicans* does not typically undergo mating, it employs several mechanisms to maintain genetic diversity, including an active parasexual cycle in which diploid cells mate and the tetraploid progeny subsequently lose chromosomes until attaining diploidy again (Bennett and Johnson 2003; Forche *et al*. 2008), frequent and extensive genetic recombination between homologous chromosomes, including multiple gene conversion events, and a remarkable tolerance of aneuploidy, which can confer selective advantage under some conditions, including increased resistance to antifungal drugs (Magee *et al*. 1992; Perepnikhatka *et al*. 1999; Hull *et al*. 2000; Magee and Magee 2000; Bennett and Johnson 2003; Selmecki *et al*. 2005; Selmecki *et al*. 2006; Coste *et al*. 2007; Rustchenko 2007; Forche *et al*. 2008; Legrand *et al*. 2008; Anderson and Bennett 2016; Mba *et al*. 2022).

*C. albicans* exhibits a number of properties associated with the ability to invade host tissue, to resist the effects of antifungal therapeutic drugs and the human immune system, and to alternately cause disease or coexist with the host as a commensal, including the ability to grow in multiple morphological forms and to switch between them, and the ability to grow as drug-resistant biofilms (Harriott and Noverr 2011; Thompson *et al*. 2011; Gow *et al*. 2012; Jacobsen *et al*. 2012; Tobudic *et al*. 2012; Mayer *et al*. 2013; Sardi *et al*. 2013; Mcmanus and Coleman 2014; Berman 2016; Legrand *et al*. 2019; Mba *et al*. 2022). Even in the commensal state in the gut and on mucosal tissues, *C. albicans* may not be as harmless as it has been assumed to be, as *Candida* interactions within the gut may set up a self-reinforcing inflammatory cycle (Kumamoto 2011; D’enfert *et al*. 2021). A recent study shows evidence of synthetically lethal infection by *C. albicans* and the bacteria *S. aureus*, in which a bacterial toxin is elicited by the fungal effect on pentose sugars (Paul *et al*. 2024). Thus, while the interplay between the fungus and the host immune system is complex, the interplay between this fungus and other members of the human microbiome may prove similarly complex.

The *Candida* Genome Database will adapt to meet these challenges so that researchers have access to as much data as possible.

### Unique aspects of CGD and the Species within it

As a knowledgebase, CGD has a number of unusual features. Foremost is the fact that the *Candida* fungi are studied mainly for their role in causing human disease, where insights into human biology–especially the immune system–come about tangentially. Said another way, *Candida* cells are not studied with the goal of understanding human cells. Within this set of pathogens, however, *C. albicans* is viewed as the model for the others because both genes and techniques have been better characterized for this species (Kabir *et al*. 2012b).

Research into *Candida* tends to take two tracks, where basic research looks into the genes and pathways that govern interactions with the host and the ability to survive under multiple stresses, while more applied research is epidemiological and clinical, i.e. looking at the response of the fungi to various drugs and/or drug candidates. While other fungi cause disease in people with weakened immune systems, e.g., species of the *Aspergillus* and *Cryptococcus* genera, these diseases are generally caused by inhalation of spores. [CDC https://www.cdc.gov/fungal/about/types-of-fungal-diseases.html] *Candida* is unique among common fungal pathogens in living as a ubiquitous commensal that becomes virulent when the immune system is compromised. Thus, study into *Candida* is especially important for understanding the “switch” between symbiotic and invasive, which often involves numerous host-pathogen interactions. For example, a recent study showed how *C. albicans* secretes a protein into host cytosol that binds and inhibits a kinase that would have begun an interferon cascade to kill the fungus (Luo *et al*. 2024). Thus, we gain insight into the human immune system when we study the ways *Candida* has evolved to evade it.

The inclusion of multiple species, as described above, also makes CGD unique. These species are more similar than different, yet have distinctive features to each. Two of the five current species (*C. glabrata* and *C. auris*) exist almost exclusively as haploid organisms, while *C. albicans, C. parapsilosis*, and *C. dubliniensis* are almost exclusively diploid. (Butler *et al*. 2009). *C. albicans, C. parapsilosis, C. auris*, and *C. tropicalis* are within the “CUG clade,” in which the CUG codon codes for serine instead of leucine (Santos and Tuite 1995; Dujon *et al*. 2004).

*C. glabrata*, as the species most closely related to *S. cerevisiae*, often acts as a “bridge” between the deep knowledge about the model yeast and the shallower knowledge about the commensal pathogenic yeasts. Like *S. cerevisiae, C. glabrata* does not make true germ tubes (Katsipoulaki *et al*. 2024) and remains in the planktonic yeast form, whereas *C*.*albicans* depends on filamentation for invasion of host tissue and loses virulence when filamentation is inhibited. (Calderone and Fonzi 2001; Witchley *et al*. 2019). The other non-albicans pathogenic species are capable of forming either pseudohyphae or true hyphae, though filamentation has varying impact on virulence (Silva *et al*. 2012; Banerjee *et al*. 2019).

Further, most pathogenic yeasts form thick biofilms composed of different cell types, including planktonic, pseudophyphal, and hyphal cells, and covered by extracellular matrix. Biofilms provide protection from immune attack and drug treatment (Sherry *et al*. 2017; De Barros *et al*. 2020). *C. glabrata*, by contrast, builds biofilms in a different manner that doesn’t rely upon filamentation. Both types of yeasts, though, use a common set of adhesins to attach to surfaces ahead of biofilm formation (Csank and Haynes 2000; Kaur *et al*. 2005; Katsipoulaki *et al*. 2024).

Among the pathogenic yeasts, *C. auris* has enhanced ability to invade skin tissues and is able to spread from skin to internal organs in a mouse model (Towns *et al*. 2024). This invasiveness on skin makes *C. auris* more problematic in nosocomial settings, where it tends to cause larger outbreaks than do the other species. Future research will almost certainly answer questions about what makes *C. auris* uniquely able to evade powerful immune responses in the skin (Ionescu *et al*. 2024; Towns *et al*. 2024).

*Candida* are exquisitely adaptive organisms, able to glean nutrients from the host while either fighting or evading the immune response. As a diploid organism with an unusually high tolerance for genomic instability, *C. albicans* is able to rapidly adapt to diverse environments (Polke *et al*. 2015). Where *C. albicans* tends to be an aggressive pathogen that battles host responses and secretes proteases to attack host cells, *C. glabrata*, in contrast, appears to rely more on stealth, trying to evade detection to aid in perseverance (Brunke and Hube 2013; Katsipoulaki *et al*. 2024). This extreme adaptability is itself a feature of these pathogens worth studying, in that the means by which these fungi adapt often tells us as much about the host as the pathogen.

## Overview of CGD content and use

### Literature Topic Tags

The *Candida* Genome Database was originally cloned from the *Saccharomyces* Genome Database in 2004 and then modified. By purposefully sharing the database organization and consistency of the user experience between the SGD and CGD knowledgebases, we allow any scientist who is familiar with SGD (of which there are a great many in the *Candida* community) to navigate the data and use the tools at CGD without requiring any extra investment of time or effort.

A weekly script imports publications into the database for annotation by a human curator. At a minimum, every publication relevant to *Candida* research gets assigned literature topic tags that provide searchability. Topic tags are assigned by reading the full paper and tagging what the results represent. Where a result pertains to a particular gene or set of genes, such as the quantification of RNA levels, the tag is assigned to those genes. Where a result pertains to the organism itself, such as the response to a drug treatment, the tag is assigned without a gene (Fig. 1).

**Figure 1.**
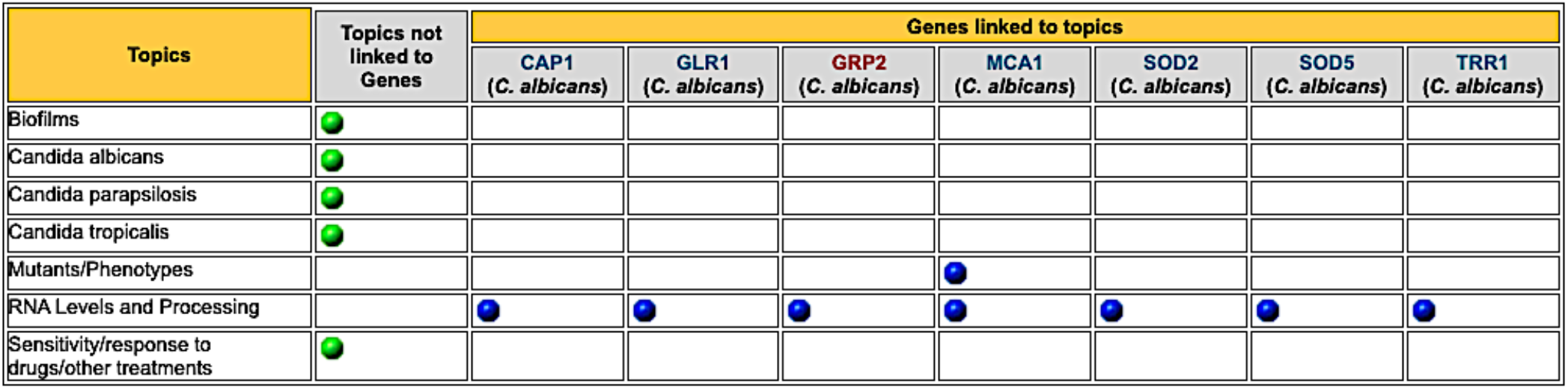
Topics addressed in (Wang *et al*. 2024). Selecting a green button will navigate to a list of other papers in CGD associated with the topic. Selecting a blue button will navigate to a list of other papers tagged for that topic for that gene.

If a researcher is interested in a particular gene, the literature associated with that gene is available with its accompanying literature tags, e.g., *GRP2* of *C. albicans* (Fig. 2). The column of “Other Genes Addressed” allows a user to identify the pathway or cluster of interest in identifying relevant literature. Similarly, an interest in mutant phenotypes can be narrowed to the relevant literature by clicking the “Mutants/Phenotypes” topic tag on the left.

**Figure 2.**
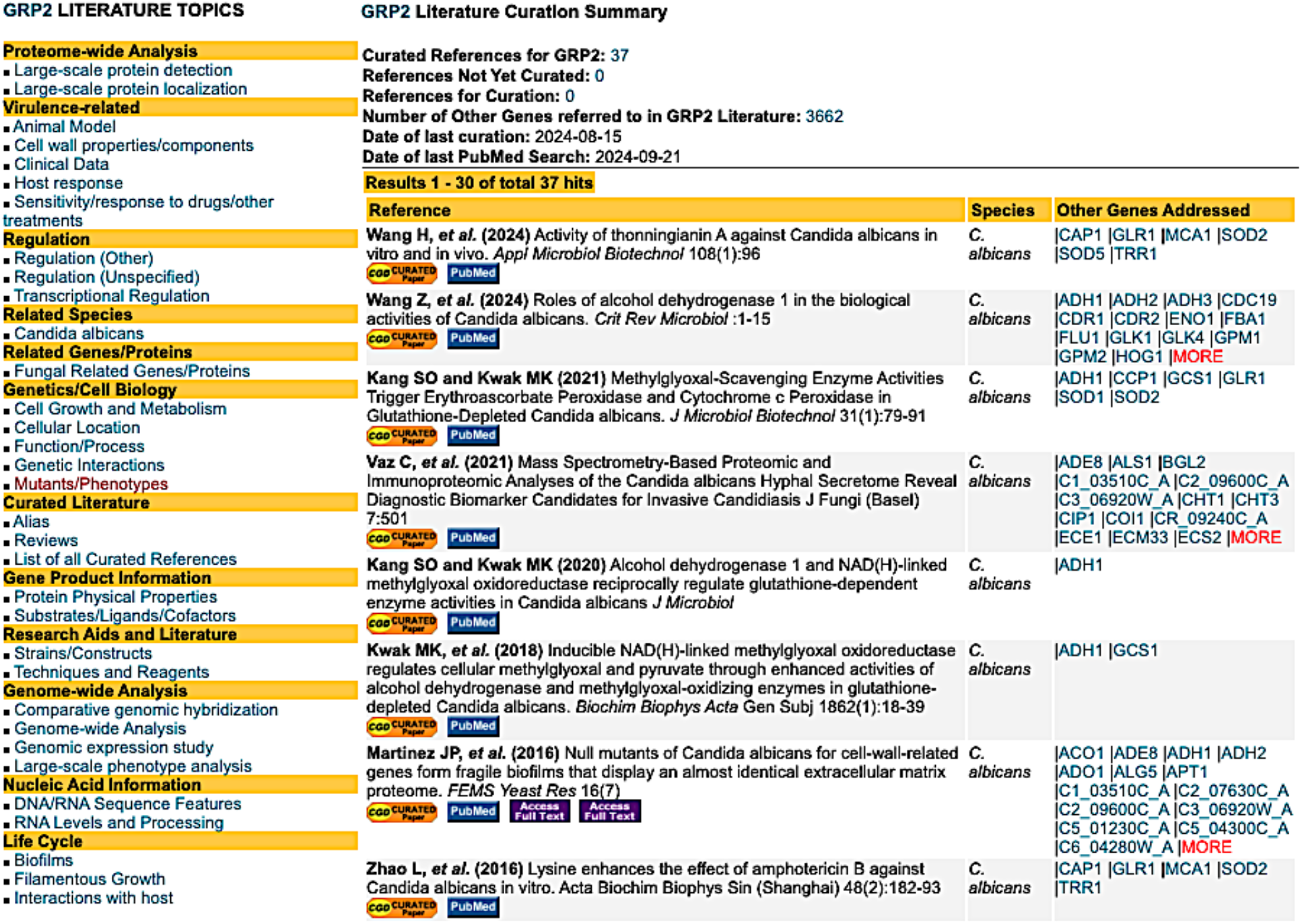
Summary of literature topics linked to curated references for *C. albicans GRP2*.

### Phenotypic Data

Beyond topic tags, CGD annotates detailed mutant phenotype data from relevant literature. Wherever a mutant is described in a publication, the observable phenotype of the mutant is annotated for the mutated or null allele. Along with the observable phenotype, the mutant is characterized for experiment type (i.e., classical genetics versus large-scale, homozygous versus heterozygous diploid versus haploid) and allele type (i.e., null versus deletion mutant or point mutation) (Fig. 3). Given the nature of many experiments on *Candida* involving drug treatment, our annotation captures the ChEBI designation for a given chemical, such that it takes only one click to see which other mutants (in all *Candida* spp. in CGD) have been characterized for response to that chemical. Also available is the strain in which the mutation was studied and any details on experimental conditions.

**Figure 3.**
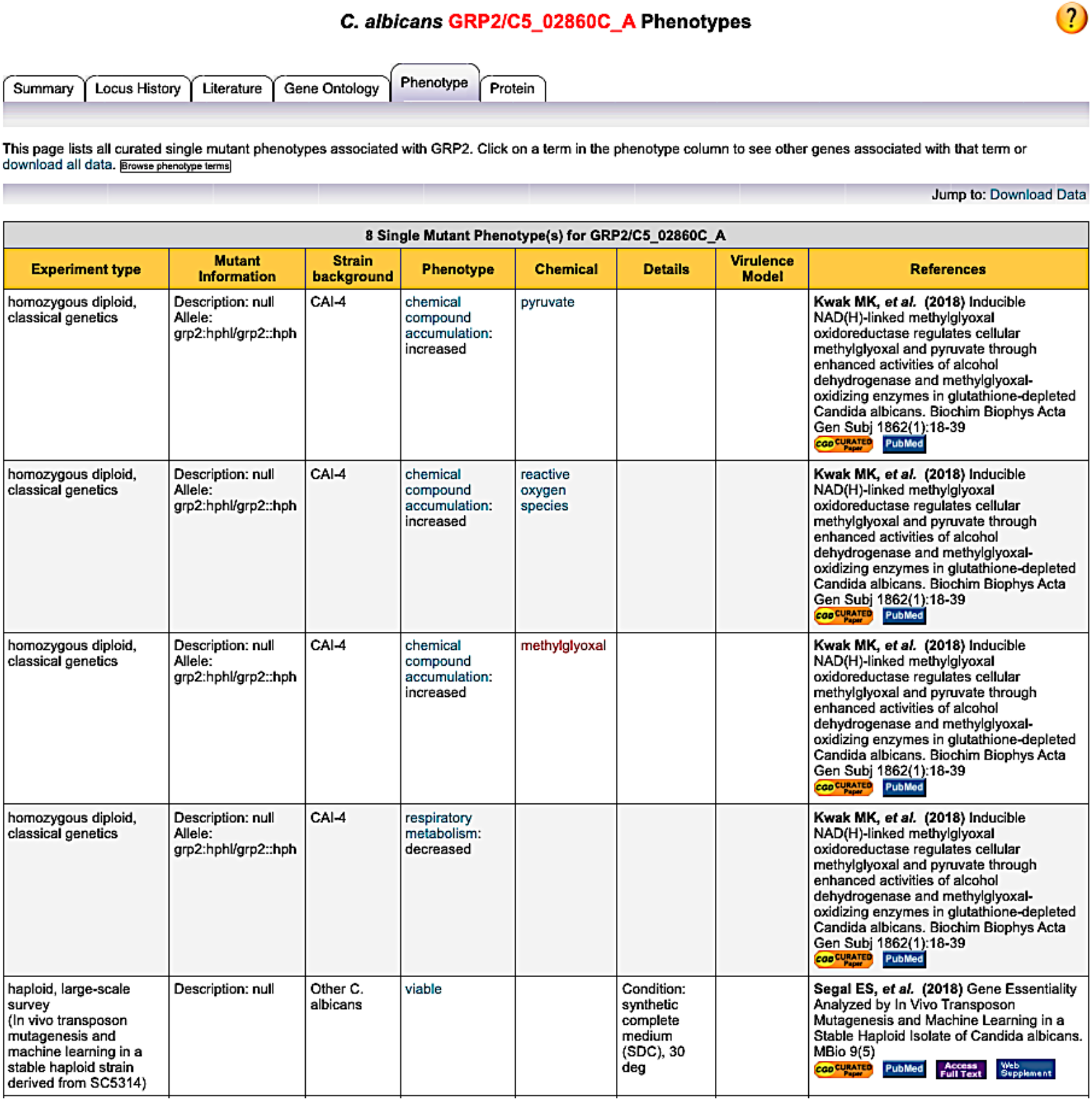
Mutant phenotype annotation for *C. albicans GRP2*.

CGD does not capture phenotypes for double or triple mutants due to the complicated nature of these interpretations. However, interactions between genes when multiply mutated are often sufficient to infer a function or process that allow designation of a Gene Ontology term.

### Gene Ontology Assignments

Manual Gene Ontology terms are assigned from publications showing experimental results supporting a given term. The terms specifically refer to the wild-type function of a given gene product; however, a mutant phenotype is often used to infer a wild-type function. When a single mutation is used to infer a term, the evidence code of Inferred by Mutant Phenotype (IMP) is applied. When multiple mutations are used to infer a role in a given function or process, it is often more appropriate to use the evidence code of Inferred by Genetic Interaction (IGI).

Inferred by Direct Assay (IDA) is the strongest level of evidence code and is preferable wherever possible as support for a term. Cellular localization terms are often supported by direct assays, as are enzyme activity molecular functions. More subtle roles in biological processes, however, often require a mutant phenotype that suggests a role, followed by direct assays designed to better test the questions.

As shown in Table 1, the number of manual GO annotations per species is directly correlated to the number of publications curated. The less-well-studied species therefore rely more heavily on computational predictions based on sequence similarity, especially Inferred from Electronic Annotation (IEA). These annotations hold more relevance where species are more closely related, such as *C. glabrata* and *S. cerevisiae*.

**Table 1.**
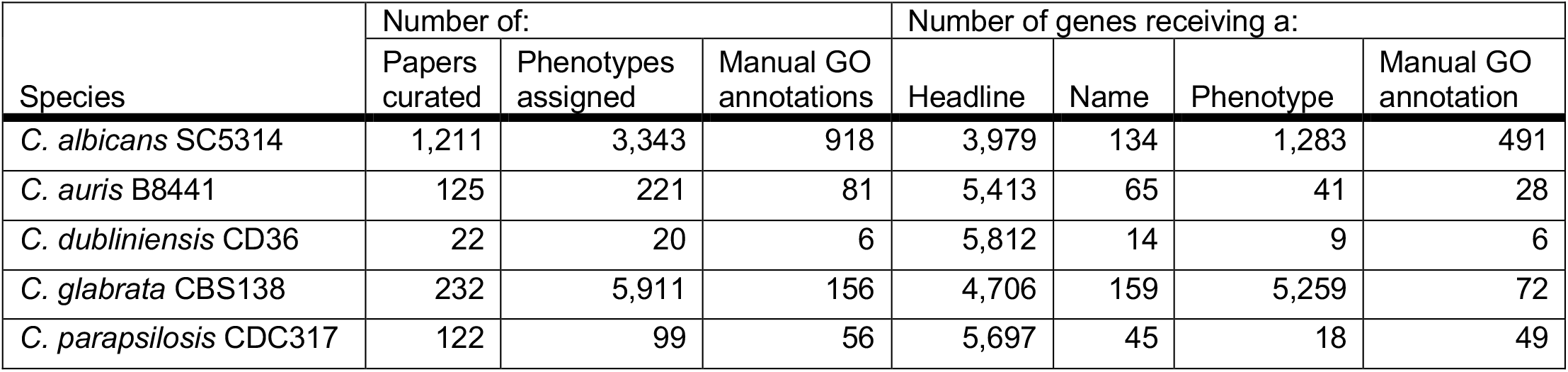
Curation statistics from 2019 to 2024.

| Species | Number of: |  |  | Number of genes receiving a: |  |  |  |
| --- | --- | --- | --- | --- | --- | --- | --- |
|  | Papers curated | Phenotypes assigned | Manual GO annotations | Headline | Name | Phenotype | Manual GO annotation |
| *C. albicans* SC5314 | 1,211 | 3,343 | 918 | 3,979 | 134 | 1,283 | 491 |
| *C. auris* B8441 | 125 | 221 | 81 | 5,413 | 65 | 41 | 28 |
| *C. dubliniensis* CD36 | 22 | 20 | 6 | 5,812 | 14 | 9 | 6 |
| *C. glabrata* CBS138 | 232 | 5,911 | 156 | 4,706 | 159 | 5,259 | 72 |
| *C. parapsilosis* CDC317 | 122 | 99 | 56 | 5,697 | 45 | 18 | 49 |

**Table 2.** Taxonomic changes to the genus *Candida*.

| Name | Synonym |
| --- | --- |
| *Candida lusitanae* | *Clavispora lusitanae* |
| *Candida krusei* | *Pichia kudriavzevii* |
| *Candida guilliermondii* | *Meyerozyma guilliermondii* |
| *Candida glabrata* | *Nakaseomyces glabratus* |
| *Candida auris* | *Candidozyma auris* |

### JBrowse Genome Browser

CGD uses the genome browser JBrowse (Buels *et al*. 2016) to present annotated features along with large-scale data in a genomic context. JBrowse allows viewing at multiple levels of resolution, from base pairs in individual sequences to broad summaries of data across large genomic regions. The display includes parallel tracks of annotated sequence features, hyperlinked to the Locus Summary pages for those features (Fig. 4). Quantitative tracks graphically display comparative information, such as relative expression level or degree of sequence conservation. JBrowse is flexible and customizable: users can easily load their own sequence datasets and analysis tracks, for display in the context of genomic features, or for comparison with datasets and tracks provided by CGD.

CGD offers a number of genomic analyses and datasets for viewing in JBrowse. For each species we provide tracks showing the level of sequence conservation at a given location, derived from whole-genome alignments between closely related species. We provide results from published gene expression experiments (RNA-seq) for *C. albicans* (8 studies as of this writing), *C. auris* (8 studies), *C. glabrata* (5 studies), *C. parapsilosis* (3 studies) and *C. dubliniensis* (2 studies). For *C. albicans* we also provide results from two chromatin occupancy experiments (ChIP-Seq), and the results from a transposon mutagenesis experiment (Segal *et al*. 2018). We also provide tracks specifying the sequence variation between two common lab strains of *C. albicans* (SC5314 and WO-1), as well as that between the two haplotypes of strain SC5314. We continually add new datasets to JBrowse as they become available, and in response to user requests.

**Figure 4.**
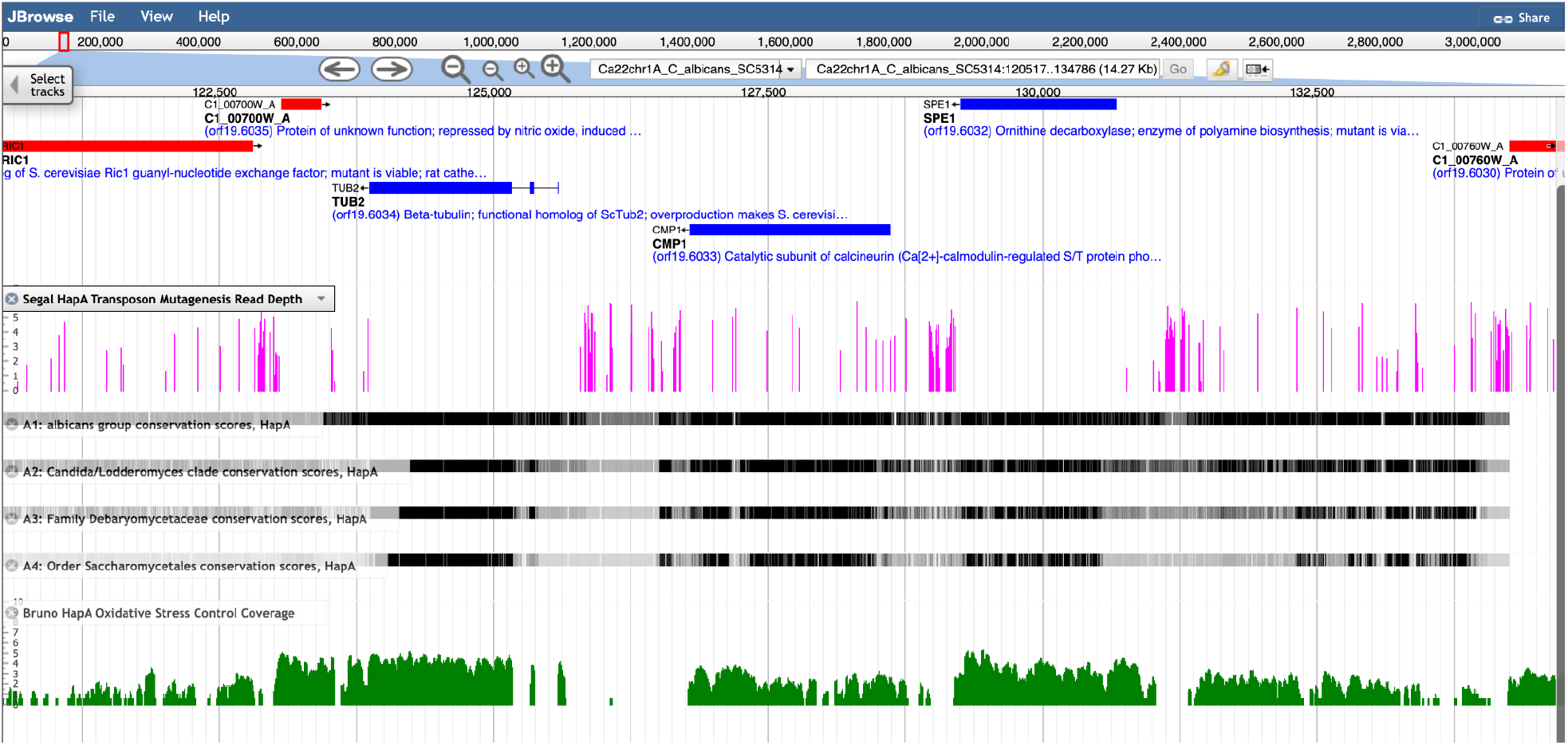
JBrowse genome browser at CGD.

A typical JBrowse display of the region around the *C. albicans* beta tubulin gene, *TUB2*. The red and blue bars in the top track of the main display window show genes annotated at CGD: red for genes encoded on the ‘W’ (plus) strand, blue for genes on the ‘C’ (minus) strand. Clicking on a bar brings up an information window for that gene and includes a link to its CGD Locus Summary Page. The vertical pink bars plot the number of hits recorded in a large-scale transposon mutagenesis (Segal *et al*. 2018) – the absence of hits in a gene indicates that the gene is essential. The horizontal black tracks show sequence conservation at four levels of divergence (the lowest track is the largest evolutionary distance); the more conserved the sequence, the more likely it is functionally important. The green bar graph shows the density of aligned reads from a typical RNA-Seq experiment (Bruno *et al*. 2010). Menus and controls at the top of the browser provide navigation, zoom and search functionalities, and allow users to load their own data.

### Downloadable Datasets

CGD strives to make as much data freely and easily available as possible. Clicking the Downloads tab brings up an explanation of the many types of downloadable data, along with links to the files (Fig. 5).

**Figure 5.**
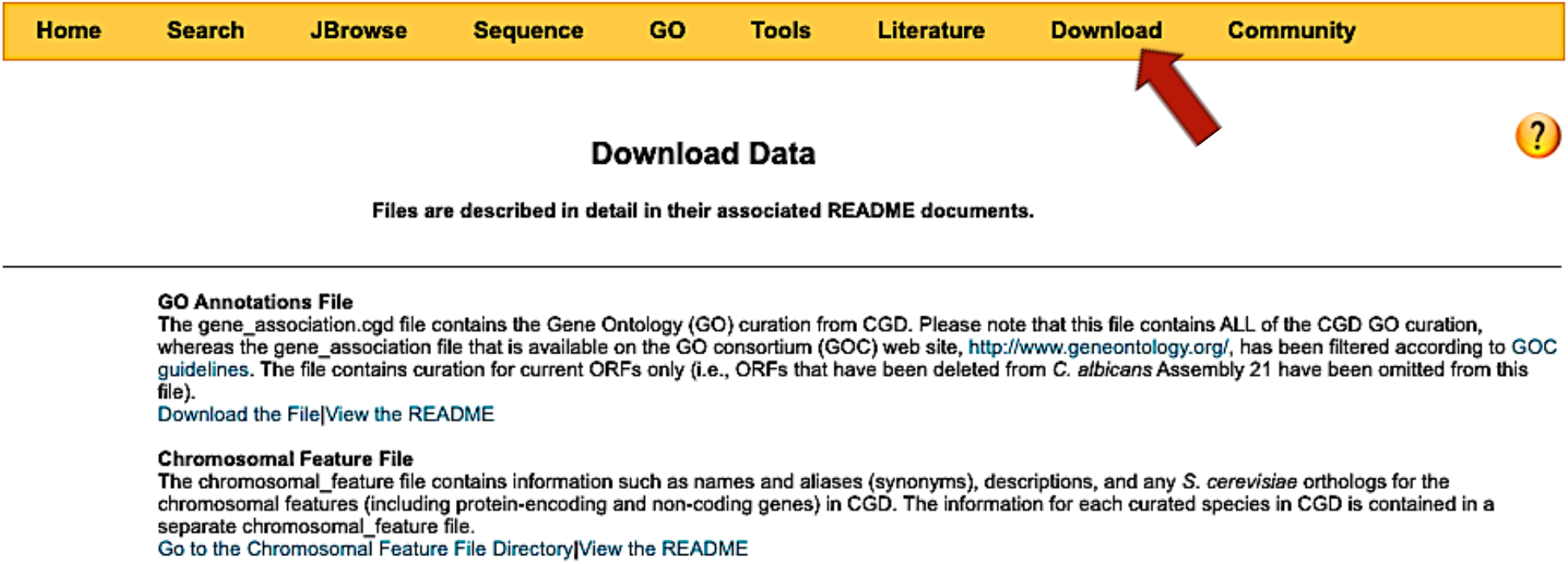
List of data files available for download by clicking the Download menu (red arrow).

Also maintained at CGD are downloadable copies of datasets from many publications (Fig. 6). We archive and provide these files to guarantee that data remains available whether or not the publication and/or data site continues to be supported online. The datasets have links to files downloadable from CGD (green buttons) and/or from the journal as a web supplement.

**Figure 6.**
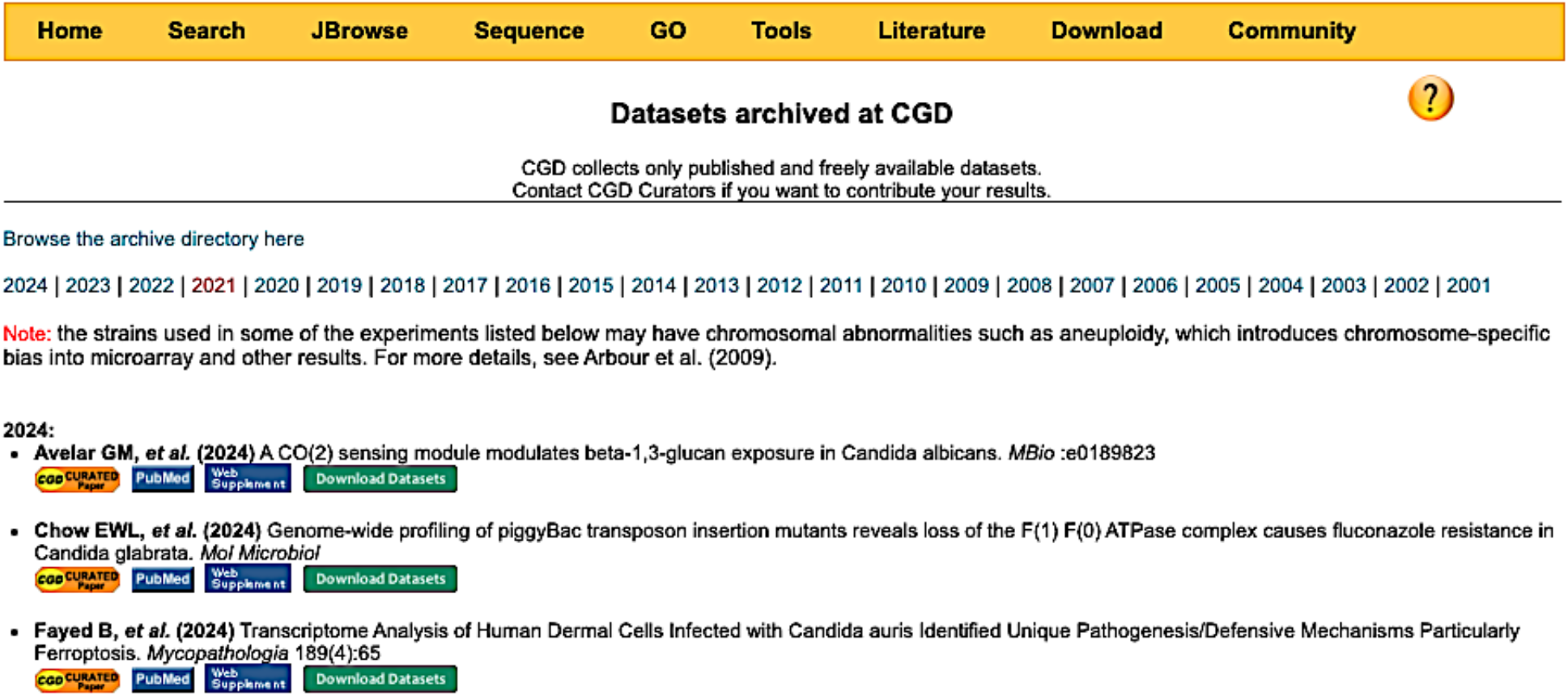
Datasets available for download on the Datasets selection of the Download menu.

### Gene nomenclature and reference genomes

The nomenclature guide resides under the Community tab (Fig. 7), as CGD incorporates names published by the *Candida* community. CGD does not assign gene names except as computational alias names based on orthology (see below).

**Figure 7.**
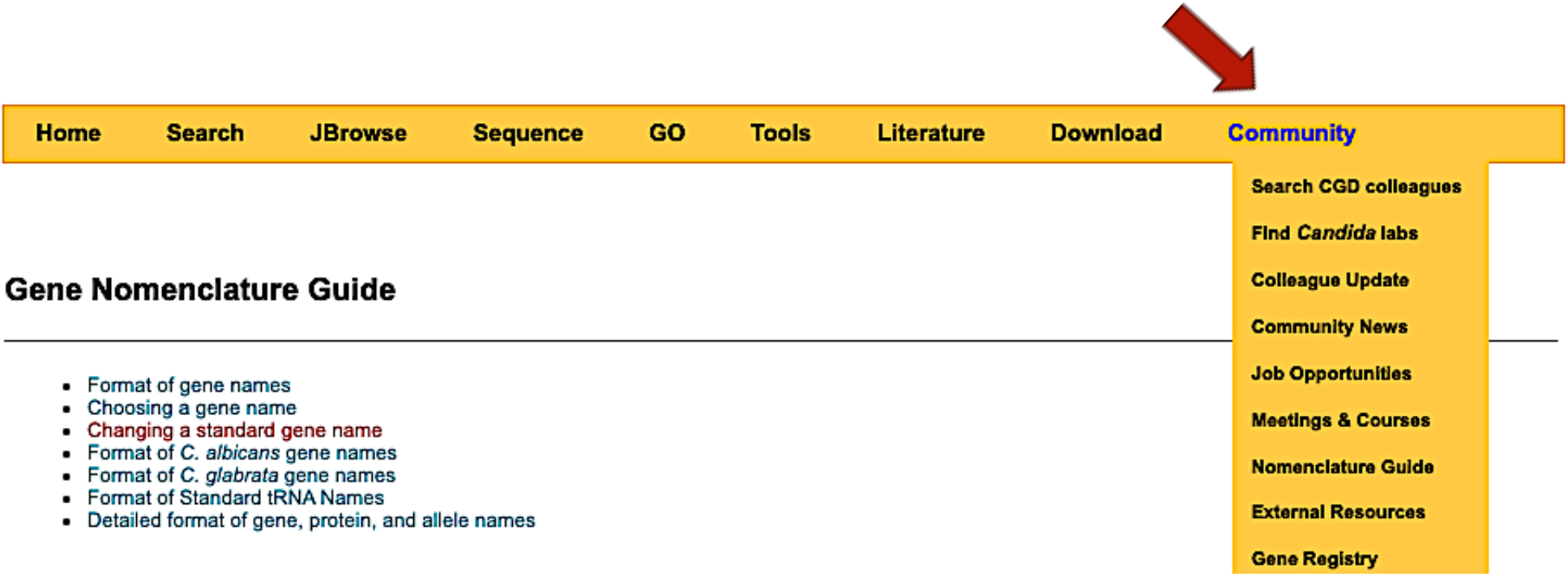
Detailed explanation of how genes are named and how names can be updated.

Standard gene names generally take the same format as for *S. cerevisiae*, with three capital letters and a number. As in SGD, exceptions to this rule are made for genes with a long-recognized non-standard name. New standard names can be reserved and then confirmed upon publication. A primary role for CGD is to moderate name changes proposed by community members, where new results suggest a name that better reflects biology. We do not make changes without consulting relevant research groups to explain the proposed changes and hopefully achieve consensus.

The maintenance of reference genomes is another key role for CGD. Genome assemblies come from multiple sources and are carefully reviewed to assess their quality. When a resequencing effort generates a new assembly of sufficient quality that changes the underlying sequence and/or sequence annotation, those changes are not only incorporated into the database, but the changes to both the sequence and the annotation are carefully recorded. We provide genome and annotation versions, so that users may download a previous, timestamped version of the genome, if for example the wish to reproduce an analysis.

Reference genomes take on a higher level of complexity in CGD because we maintain them for five organisms, some of which are typically diploid and others typically haploid. Further, *C. albicans* has A and B haplotypes that differ for some genes and not others (Fig. 8). Every *C. albicans* locus page thus has a field for “Allelic variation,” where we note “No allelic variation in feature” versus “Non-synonymous variation between alleles.”

**Figure 8.**
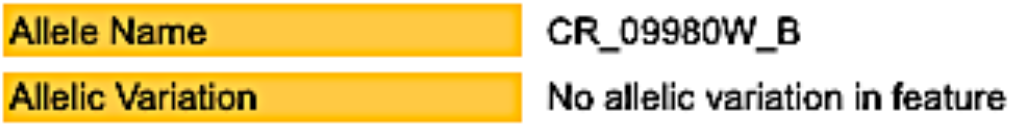
Example of haplotype information for a *C. albicans* gene.

For users interested in allelic variation, each feature has a link to the JBrowse page that allows you to view either haplotype (Fig. 9). We also align genomic expression datasets to each *C. albicans* haplotype separately, so that expression can be viewed for either allele or set of alleles.

**Figure 9.**
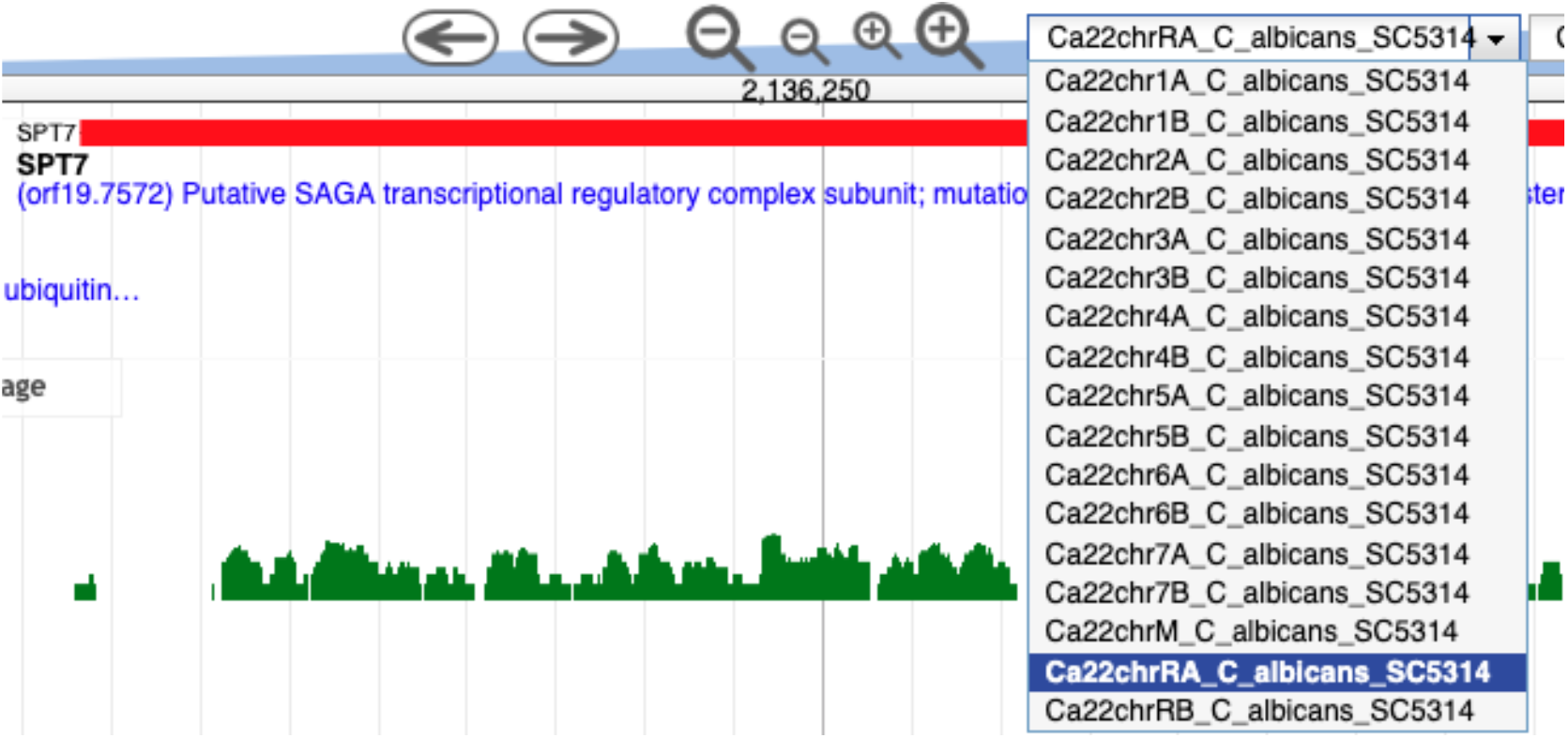
Example of expression data available by haplotype for *C. albicans SPT7*.

Sequence data are fully retrievable from our Downloads menu, along with information on sources, software for unzipping files, and available formats. We maintain sequence for older assemblies as well as current assemblies, making each separately accessible.

## Orthology Resources

Orthologs are genes in different organisms that descended from a common ancestral gene (Koonin 2005). The assumption that the orthologs of a gene with known function will share a similar function has proven to be a powerful concept in evolutionary genomics, allowing for the annotation of many if not most of the genes in newly sequenced genomes (Dessimoz *et al*. 2012). While annotations based solely on orthology predictions should be considered provisional pending experimental confirmation–as ortholog functions could have diverged in organisms living in different environments or the predictions themselves could be faulty– ortholog assignments provide researchers with an excellent starting point for focusing their functional studies.

At CGD we use the ortholog assignments for *Candida*-related species provided by the *Candida* Gene Order Browser (CGOB, http://cgob3.ucd.ie/cgob.pl; (Maguire *et al*. 2013)). The ortholog predictions of CGOB are particularly powerful because they are based on gene synteny – conserved chromosomal gene positions among closely related organisms – in addition to sequence homology, thus increasing the confidence that the assigned genes are related by direct descent. CGD provides the following resources for *Candida* orthologs:

### Ortholog Downloads

For each pair of CGD-curated species, we provide a downloadable list of orthologs between those species: http://www.candidagenome.org/download/homology/orthologs/. For genes that have no ortholog assignment, we provide lists of best BLAST hits between species: http://www.candidagenome.org/download/homology/best_hits/.

### Ortholog section of the Locus Page

On the Locus Page of each gene at CGD, we provide links to the orthologous gene in each of the other CGD-curated *Candida* species, as well as a link to the CGOB page for that group of orthologs. If available, we also link to orthologous genes in non-*Candida* species, such as *S. cerevisiae* and *S. pombe*.

### Homologs tab of the Locus Page

One of the tabs at the top of each gene’s Locus Page is for Homologs of that gene. The Homologs tab provides these additional resources:

1. An expanded list of orthologs in other *Candida*-related species, including species not yet curated by CGD, plus downloadable sequence files for those orthologs;
2. Phylogenetic trees showing the degree of relatedness among the orthologs, plus downloadable trees in formats readable by most bioinformatics programs;
3. Multiple sequence alignments for both protein and coding sequences, highlighting conserved regions among the orthologs, plus downloadable alignments in ClustalW and FASTA formats.

## Interactions with community

### Diverse colleagues

CGD makes an effort to serve as a central hub for a diverse set of research groups. Because *Candida* is studied as a pathogen as well as a model, some groups focus on basic research into *Candida* biology while many others perform clinical studies. The clinical studies include epidemiology and phylogeny of clinical isolates that have varying degrees of virulence and drug resistance.

Several of the most virulent Candida species, including *C. glabrata* and *C. auris*, possess inherent resistance to the most common antifungals. Thus, many researchers seek new antifungals from across a broad spectrum of organisms. Others seek new drugs in chemistry. The diversity of these research interests makes it important that CGD provides services to all groups.

Our Community tab includes a list of laboratories and colleagues to facilitate connections among groups. The basic research into cellular biology of the fungi will undoubtedly inform the work of those seeking improved clinical tools.

### Quarterly newsletter

CGD has instituted a quarterly newsletter for which the goal is to disseminate timely news pertinent to the efforts of many research groups. We highlight improvements or changes to the database itself, but also highlight upcoming meetings and other relevant announcements.

Each newsletter includes a Research Spotlight, in which we describe a novel recent finding in abbreviated format. The spotlight serves to disseminate news of important progress as well as foster a sense of community.

### Moderation

Within a research community, the aid of a neutral party in moderating communications is often of great value. CGD performs this role in several ways, but primarily in the realm of proposed changes to gene names. We work with the research group suggesting a change to develop the proposal, and then contact the research groups with a stake in the previous nomenclature. Having the proposal come from a neutral party often facilitates the exchange.

Another area of moderation is in regard to taxonomic changes within the set of pathogens covered by the database. Further genomic studies into the *Candida* genus have made clear that several of the current species do not belong in the genus due to extended evolutionary distances. The community as a whole will likely adopt these changes; however, the timing of the adoption can either minimize or maximize confusion. CGD hopes to assist in minimizing the impact.

### Community Surveys

Every five years, CGD undertakes a community survey to gauge the changing needs of the community. The survey asks how the database is currently used and how it could better serve the need. Further, because CGD incorporates multiple species of *Candida*, we ask the community for input on which species to next consider for a full set of locus pages.

## Future Challenges

### Navigation of many expression datasets

As the techniques for transcriptomics become easier to apply, assessment of genome-wide gene expression has become a standard tool for *Candida*. As a knowledgebase, we balance the need to make these datasets accessible with the realization that the wealth of data is becoming overwhelming. Where it was once possible to align raw datasets to our reference genomes for every study, there are now too many for this to remain feasible. We thus prioritize studies of general interest over those focusing on a narrow question, e.g. the effect of knocking out a single gene.

The next logical step is to collate data in software tools that illustrate the patterns within metadata. CGD has one such tool that was developed for illustrating microarray data. We are currently working toward adapting this tool for more recent forms of genomic expression data, such as RNA-seq. The eventual output will be a heatmap that links genes to datasets.

### Navigating and understanding multiple genomes

With the increasing incidence of drug-resistant *Candida* infections proving lethal in hospital settings, researchers are conducting greater numbers of epidemiological studies across the globe. What they’re finding is a greater number of different *Candida* species causing virulent infections than was originally believed to be the case.

For the CGD knowledgebase, this means the addition of multiple species over time. The most recent addition was *C. auris*, with locus pages for every feature and multiple reference genomes against which users can BLAST query sequences. These pages have been filling quickly with experimental evidence generated by a suite of new researchers studying this deadly and highly transmissible pathogen.

Our community survey of 2024 identified *Candida tropicalis* as the next species for which the community would like full coverage by CGD. The addition of this species is one of our future goals; however, the research coverage for this species does not yet warrant locus pages for genomic features that would largely remain uncharacterized into the foreseeable future. We will, of course, provide the reference genome for searching and downloading as needed.

### How to integrate artificial intelligence

The rise of artificial intelligence has knowledgebases asking how to integrate this powerful technology. From the perspective of those maintaining a database, there are numerous possibilities for using AI to perform tasks that are currently performed manually. Such tasks include reading abstracts to triage literature for input to the database, writing summary paragraphs from sets of annotations, or comparing orthologous genes to glean commonalities that might have biological relevance.

Alternatively, from the user perspective, AI could be used to mine the data in the database in ways we cannot currently support. For example, AI might allow a user to make a request such as “List all the genes upregulated during oxidative stress that are also upregulated during hyphal growth.” From this set, the user might ask a further question such as “Do any studies identify members of this set of genes as important for drug resistance?”

We will integrate AI in a measured strategy that prioritizes straightforward applications and prepares for more complicated usage. As an example of a straightforward application, the advent of large language models (LLMs) that employ natural language are especially suited to summarizing compiled annotation and returning human-sounding summary text. CGD does not have the available staff to develop such tools itself, but such tools are being developed by the Alliance of Genome Resources Consortium. We will keep apprised of the toolset they develop and, when appropriate, deploy such tools to CGD.

